# Rapid Identity and Quantity CQA Test for Multivalent mRNA Drug Product Formulations

**DOI:** 10.1101/2022.08.31.506088

**Authors:** Rachel Y. Gao, Christine M. Riley, Evan Toth, Rebecca H. Blair, Megan N. Gerold, Caitlin McCormick, Amber W. Taylor, Tianjing Hu, Kathy L. Rowlen, Erica D. Dawson

## Abstract

The COVID-19 pandemic highlighted mRNA as a promising platform for vaccines and therapeutics. Many of the analytical tools used to characterize the critical quality attributes of mRNA are inherently singleplex and are not necessarily optimal from a labor and cost perspective. Here we demonstrate feasibility of a multiplexed platform (VaxArray) for efficient identity verification and concentration determination for both monovalent and multivalent mRNA formulations. A model system comprised of mRNA constructs for influenza hemagglutinin and neuraminidase was used to characterize the analytical performance metrics for a VaxArray mRNA assay. The assay presented herein had a time to result of less than 2 hours, required no PCR-based amplification nor extraction of mRNA from lipid nanoparticles, and exhibited high construct specificity that enabled application to the bivalent mixture. The sensitivity for influenza hemagglutinin and neuraminidase mRNA was sub-µg/mL, which is vaccine-relevant, and the average accuracy (%recovery) and precision were 104%±2%and 9%±2%, respectively.

## INTRODUCTION

The COVID-19 mRNA vaccines developed in rapid response to the March 2020 global pandemic clearly demonstrated the capabilities of this now critical vaccine platform.^1-12^ Given the proven safety and efficacy of the mRNA-based COVID-19 vaccines that have received emergency use authorization or licensure to date,^13,14^ mRNA vaccines will continue to be important to global public health moving forward. This supposition is supported by the numerous vaccines currently in pre-clinical and clinical testing.^10,11,15-17^ One significant benefit of mRNA-based vaccines over traditional protein-based vaccines is the speed and straightforward scalability of the manufacturing process,^11,17-20^ highlighted by the authorization of both the Pfizer/BioNTech and Moderna mRNA vaccines only 9 months after the COVID-19 pandemic was declared. While developers and manufacturers are not likely to keep up a pandemic pace for all vaccines, the timeline for availability of future mRNA vaccines is anticipated to be significantly shorter than the typical time for traditional vaccine development time of 15 years or more.^4,21^

While the initial COVID-19 vaccines on the market in the US are monovalent, future mRNA vaccines are likely to be increasingly multivalent. Some examples are Moderna’s bivalent COVID-19 vaccines, mRNA-1273.214 and mRNA-1273.222, each containing two unique COVID-19 directed mRNA constructs, both currently in Phase 3 trials. ^22-24^ In addition, Pfizer/BioNTech has a bivalent COVID-19 vaccine, targeting both the BA.4 and BA.5 variants, likely to be available in Europe in Fall 2022.^25^ Another high priority for multivalent mRNA vaccines is influenza, with candidates including Moderna’s mRNA-1010 quadrivalent formulation,^26,27^ Pfizer’s bivalent influenza modRNA vaccine (bIRV),^28,29^ and GSK/CureVac’s CVSQIV^30^ currently in clinical trials. In addition, combinations of respiratory viruses in a multivalent mRNA vaccine (*i*.*e*. coronavirus and influenza, influenza and RSV) are also in development or likely on the horizon, such as Moderna’s combined COVID-19/ quadrivalent influenza candidate (mRNA-1273 and mRNA-1010).^31^

A variety of methods are recommended for identity and quantification of mRNA, as outlined in the recent draft USP guidance document for mRNA vaccine analytics, in which industry input and discussion were encouraged.^32^ In the draft USP guidance, sequencing and reverse transcriptase polymerase chain reaction (RT-PCR)-based methods are outlined to confirm mRNA identity.^32^ While sequencing provides high information content, the upfront mRNA enrichment, isolation, amplification, ligation, and other sample processing and data analysis steps significantly increase complexity, time to result, and cost which may limit adoption by lower- and middle-income country vaccine manufacturers.^33^ While more straightforward than sequencing, RT-PCR based identity testing still requires special workflows and sample handling to eliminate potential for cross-contamination with amplifiable nucleic acids and requires upfront purification if the mRNA is encapsulated in a lipid nanoparticle (LNP).^32,33^

Recommended methods for mRNA quantitation are PCR-based methods and UV spectroscopy. UV spectroscopy is straightforward on relatively pure bulk mRNA, and the UV-based RiboGreen assay can be utilized to quantify mRNA against a standard curve on LNP-encapsulated materials. However, UV-based methods suffer from interference in unpurified preparations and importantly lack the specificity required for simultaneous individual quantification of multiple mRNA constructs in multivalent samples. While digital PCR can quantify multiple mRNA sequences, it requires upfront mRNA extraction from complex matrices such as LNP-encapsulated samples, with extraction processes requiring up to a day prior to analysis.

These current methodologies were adopted out of the urgent need for COVID-19 vaccines to address the pandemic, and the speed of development was laudable.^11,17,20,33^ While future mRNA vaccines may not be developed at quite the same pace as those for COVID-19, anticipated accelerated development timelines compared to traditional vaccines and an increase in development of *multivalent* mRNA vaccines highlight the need for improved, rapid analytical tools that can streamline bioprocess development and optimization, formulation development, and QC testing to further reduce time to market and improve between-lab standardization.^21,32,34^ In this work, we demonstrate performance of a model multiplexed nucleic acid microarray-based assay for influenza mRNA vaccine construct identity and quantification. Performance is demonstrated using vaccine-relevant model influenza hemagglutinin (HA) and neuraminidase (NA) bivalent mRNA constructs from the literature.^35^ The assay is based on a modification of InDevR’s VaxArray immunoassay platform for antigen quantification in traditional protein-based vaccines,^36-38^ currently in use by a wide variety of vaccine developers and manufacturers worldwide. The VaxArray influenza mRNA assay presented herein is rapid (less than 2 hours), requires no mRNA extraction or purification (even in LNP-encapsulated samples) or RT-PCR amplification, has a variety of flexible capture and detection schemes for different applications, and provides identification, simultaneous quantification of multiple constructs in a bivalent mixture with high accuracy and precision, and vaccine-relevant limits of quantification. Here, we describe this assay and highlight relevant performance data in both naked mRNA (that is, with no LNP components present) and LNP-encapsulated mRNA.

## RESULTS

### VaxArray mRNA assay design

As described in the Methods section, 37 sequences were designed to target the coding regions of the NA and HA mRNA constructs (21 for NA and 16 for HA) and were used either as microarray capture oligonucleotides (oligos) or detection labels. All capture oligos were initially tested to assess basic reactivity and target specificity for suitability of inclusion in the final assay (data not shown). Eight capture oligos were chosen (four targeting each mRNA construct), as schematically shown in **Figure 1. Figures 1a** and **1b** show the general structure of the NA and HA constructs, respectively, along with the associated capture oligos. Performance data herein focus on two coding region capture oligos for each construct, highlighted in orange. A 30-mer polyT capture oligo targeting the 3’ poly A tail was also included on the microarray. Each microarray slide contains 16 replicate arrays as depicted in **Figure 1c**, with the final microarray layout (each capture oligo printed in 9 replicates) shown in **Figure 1d**. This array design allows flexibility in the capture and detection labeling of mRNAs, allowing either the capture or labeling step to be construct-specific by targeting the coding region, or universal by targeting the polyA tail or 5’ cap to address different applications. Some possible detection modes are shown schematically in **Figure 1e**. Most data to be highlighted here focused on capturing the mRNA in the respective coding region and detection via a polyT oligo detection label, as shown in **Figure 1e(i)**. Alternatively, labeling can be performed with an anti-5’ cap antibody, as shown schematically in **Figures 1e(ii)** and **1e(iii)**, with **1e(iii)** enabling detection of full-length mRNA (from tail to cap) in monovalent formulations. Alternatively, the coding region oligonucleotides designed can be used as fluorescent labels to enable detection via the coding region as shown in **Figures 1e(iv) and 1e(v)**.

**Figure 1.**
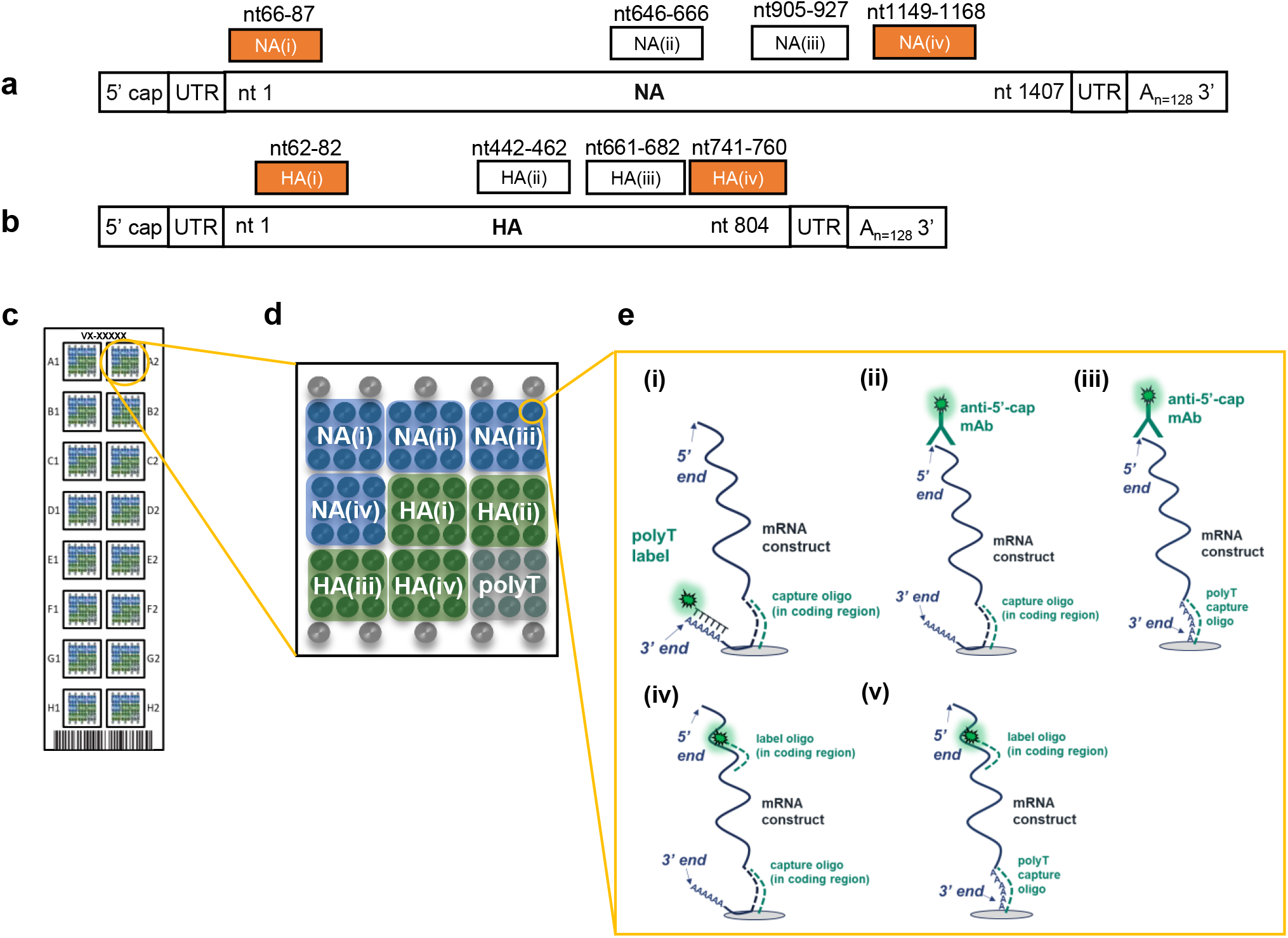
(a and b) Schematic of the HA and NA constructs used, respectively, with binding location of oligonucleotide captures shown above each construct. Captures shown in orange were the focus of performance assessment (c) Schematic representation of the VaxArray mRNA microarray slide showing 16 replicate microarrays, (d) individual microarray layout showing 9 replicate spots for each capture oligo, and fiducial markers in grey in top and bottom rows, (e) assay detection schemes represented herein, namely (i) coding region capture and polyT labeling, (ii) coding region capture and 5’ cap labeling, (iii) polyT capture and 5’ cap labeling, (iv) coding region capture and coding region labeling, and (v) polyT capture and coding region labeling.

Briefly, the general assay protocol (described in detail in the Methods section) takes less than 2 hours and consists of adding the mRNA(s) of interest to the chip and incubating in an optimized buffer to allow hybridization, washing, adding the detection label of interest and incubating, followed by washing and fluorescence imaging with the VaxArray Imaging System. Quantitative image analysis is automatic with the associated 21CFR11 software.

### Assay shows high specificity and reactivity for HA and NA targets

**Figure 2a** shows the microarray layout alongside representative fluorescence images of monovalent NA, HA, and bivalent NA/HA, all analyzed at 10 µg/mL mRNA and detected via the polyT oligo. These images qualitatively indicate that the microarray capture oligos designed for each mRNA construct generate specific signal for the intended target. Quantitative analysis of the resulting pixel intensities indicates signals vary for each oligo designed, likely due to different binding affinities. Capture oligos designed for HA mRNA constructs were highly specific, as indicated by no resulting signal above background in the presence of NA mRNA. Capture oligos designed for detection of NA were equally specific. These data are shown in the highlighted rows of **Figure 2b** in which the monovalent samples generated signal to background ratios (S/B) on the target capture oligos ranging from 8.1 to 26.6, indicating strong reactivity.

**Figure 2.**
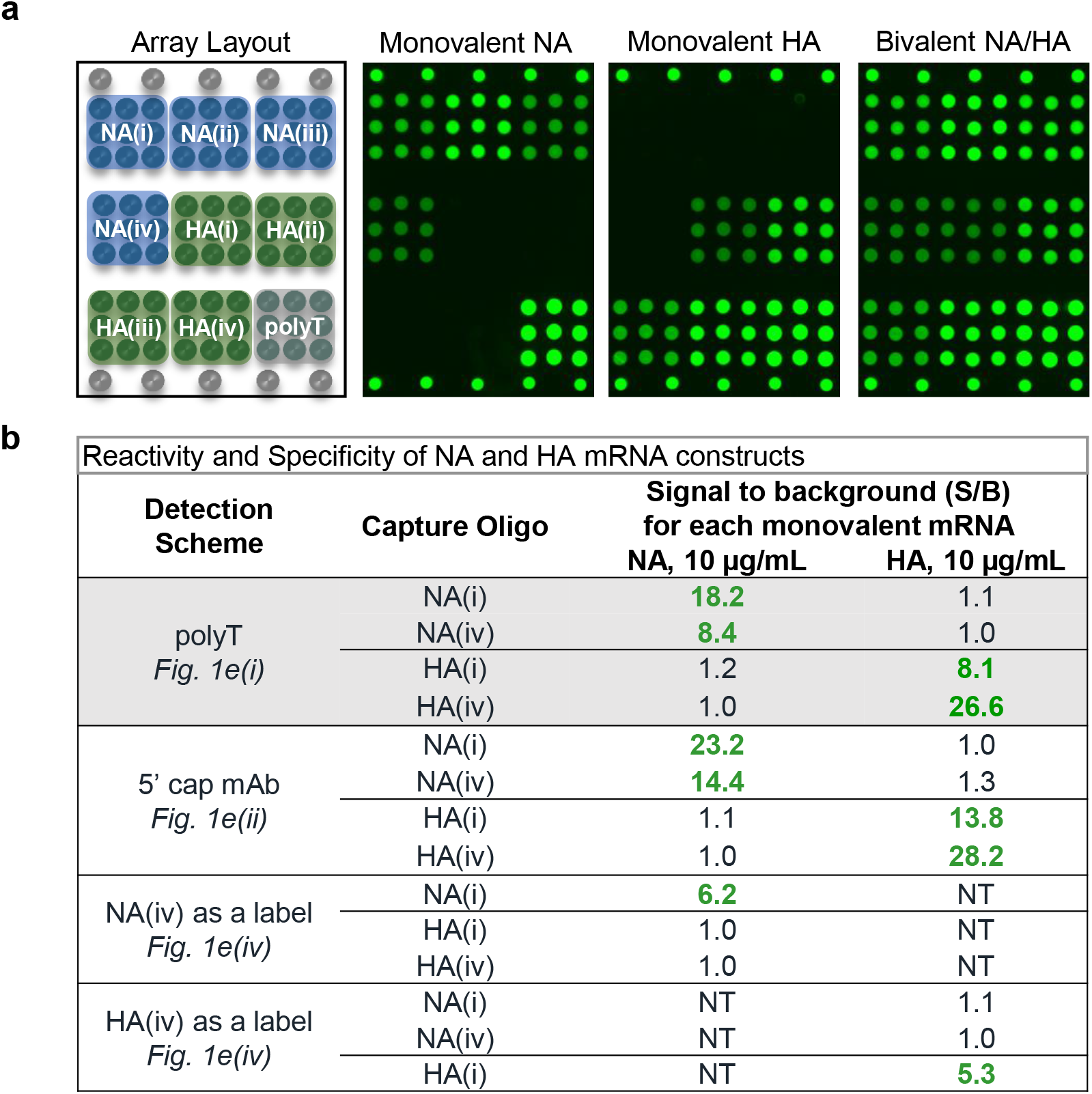
Naked mRNA reactivity and specificity. (a) Representative microarray images demonstrating reactivity and specificity of 10 µg/mL mRNA using the polyT label. (b) Signal/background (S/B) for monovalent NA and HA for four different labeling schemes (average of n=3 for each value). Text in bold green shows S/B >3 generated, which indicates positive signal. NT indicates not tested.

Reactivity and specificity for a variety of detection schemes including coding region capture/anti-5’ cap labeling (**Figure 1e(ii)**) and coding region capture/coding region labeling (**Figure 1e(iv)**) were also investigated. In all cases, good reactivity, as indicated by S/B values ranging from 5.3 to 28.2 for the intended construct, and good specificity, as indicated by S/B ratios of ≤ 1.3 for the off-target construct as shown in **Figure 2b**. These data indicate highly specific detection of multiple mRNAs in monovalent and multivalent samples for a variety of assay detection principles, with demonstrated specificity in monovalent samples important for use as an identity test.

### Monovalent mRNA samples show excellent linear response, and bivalent response matches that of monovalent

**Figure 3** highlights the average response for triplicate 8-point response curves on the NA(i), HA(i), and polyT capture oligos for monovalent NA (left column) and HA (right column) mRNA using four different capture and labeling strategies. All four detection schemes shown, represented by **Figures 1e(i-iv)** demonstrate good linearity with dilution, with a single linear fit R^2^ > 0.96. Corresponding data for capture oligos NA(iv) and HA(iv) can be found in **Supplementary Figure 1**. Data in **Figure 3a** shows good linearity with response for coding region capture and universal labeling via the polyA tail over a vaccine-relevant concentration range, important for application to mRNA quantification in bioprocess development and optimization. **Figure 3b** highlights response curves for capture via the 3’ polyA tail and labeling via the 5’ cap using an anti-5’ cap antibody at the opposite end of the mRNA construct. This detection scheme enables quantification of full-length mRNA in monovalent samples as a measure mRNA integrity. **Figure 3c** highlights linearity of response for capture in the coding region for each construct and subsequent universal labeling of the 5’ cap using an anti-5’ cap antibody, demonstrating viability of another universal labeling approach. **Figure 3d** demonstrates the ability for the assay to capture in the coding region and be labeled in the coding region for added specificity given that the labels are also specific to each construct, adding additional potential benefit for identity testing by targeting more than one portion of the coding region.

**Figure 3.**
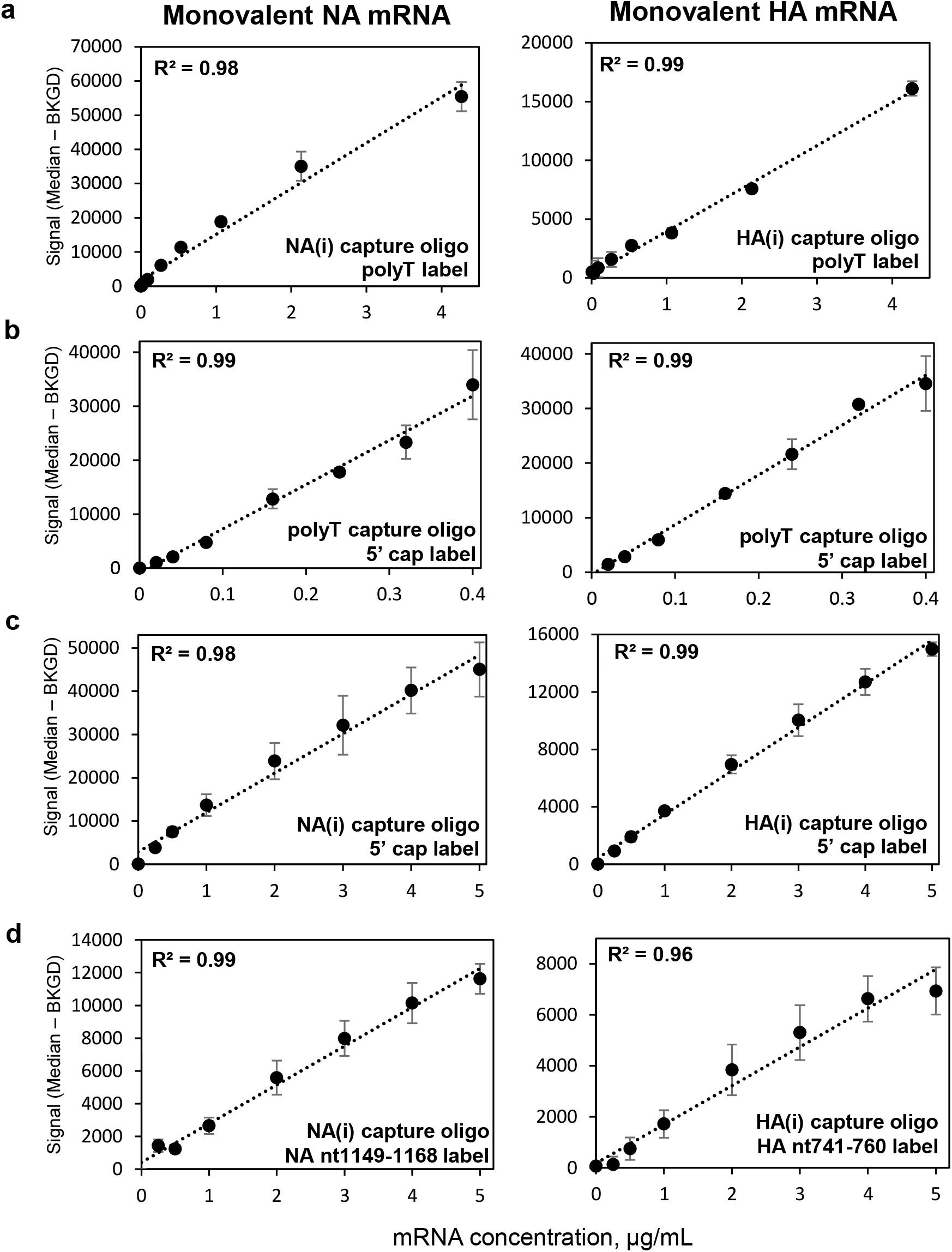
Linearity of response for a variety of capture and detection schemes in monovalent mRNA samples, with NA in left column and HA in right column. (a) Coding region capture with polyT label as shown in Figure 1e(i), (b) polyT oligo capture with 5’ cap antibody label as shown in Figure 1e(iii), (c) Coding region capture and labeling with 5’ cap antibody label as shown in Figure 1e(ii), (d) coding region capture with coding region labeling as shown in Figure 1e(iv). Data points are the average of three replicates and error bars indicate ± 1 standard deviation. R^2^ in upper left of each plot is based on a single linear regression.

**Figure 4.** shows a comparison of VaxArray signal responses for each monovalent mRNA on its respective capture oligos as well as a bivalent mixture of NA/HA mRNA at equal concentrations of each mRNA, all labeled with the polyT oligo detection label. As shown in **Figures 4a** and **4b**, the response curves for the NA mRNA component in a monovalent vs. bivalent preparation in which HA mRNA was also present are quite similar. Likewise, in **Figures 4c** and **4d**, the response curves for the HA mRNA component are also quite similar, regardless of whether alone or in a bivalent mixture. Importantly, these data show no interference from the off-target mRNA in the presence of the target mRNA and indicate independent quantification is feasible in a bivalent mixture. Given the similarity of responses in **Figure 4**, other analytical performance data presented herein were assessed based on bivalent HA/NA mixtures.

**Figure 4.**
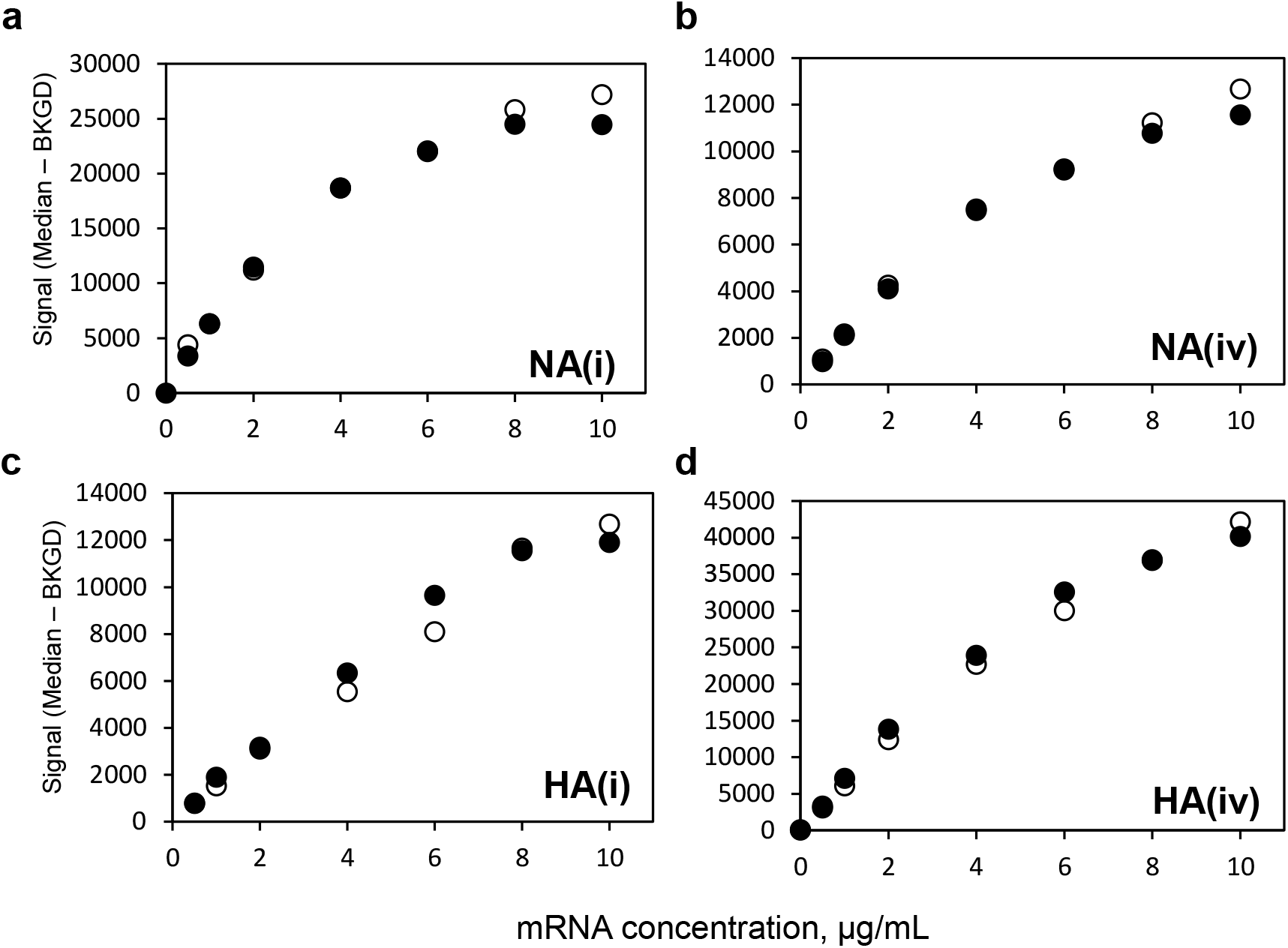
Similarity of response for monovalent (•) and bivalent (○) 8-point dilution series using polyT oligo detection label for NA mRNA using (a) NA(i) coding region capture, (b) NA(iv) coding region capture, and for HA mRNA using (c) HA(i) coding region capture, and (d) HA(iv) coding region capture.

### Assay has vaccine-relevant sensitivity and at least ∼100x working range

Fifteen (15)-point curves of bivalent NA/HA, at equal concentrations in the mixture, were analyzed to approximate the lower and upper limits of quantification, LLOQ and ULOQ, respectively, for both mRNA constructs (data not shown) for coding region capture and polyT labeling. The approximate LLOQ was determined as the concentration at which the signal generated was equal to the average blank signal + 5σ (where σ = standard deviation of the background signal). The ULOQ was approximated as the concentration at which the signal was 90% of the maximum observed signal. After determining the approximate limits, the LLOQ and ULOQ were verified by determining the lowest and highest concentrations that generated appropriate precision and accuracy as defined as RSD <15% and 100 ± 15% recovery for four replicates analyzed at concentrations near the approximate LLOQ and ULOQ. **Table 1** shows the values obtained, with LLOQ ranging from 0.08 μg/mL to 0.2 μg/mL, and ULOQ ranging from 14 μg/mL to 18 μg/mL. The dynamic range (ULOQ/LLOQ) for all four coding region capture oligos was > 90x. Given mRNA construct concentrations of 200 µg/mL in on market COVID-19 vaccines and a pandemic influenza mRNA vaccine in a phase 1 study,^21-23^ these data demonstrate an ability to quantify over a vaccine-relevant concentration range for application to formulated vaccine samples. For bioprocess samples (bulk mRNAs) that are likely at significantly higher concentration, a larger upfront dilution can easily bring samples into the appropriate working range.

**Table 1.** Analytical Sensitivity and Range

| **Capture Oligo** | **LLOQ (µg/mL)** | **ULOQ (µg/mL)** | **Linear Dynamic Range (ULOQ/LLOQ)** |
| --- | --- | --- | --- |
| **NA(i)** | 0.08 | 14 | 175 |
| **NA(iv)** | 0.20 | 18 | 90 |
| **HA(i)** | 0.20 | 18 | 90 |
| **HA(iv)** | 0.08 | 17 | 213 |

Bivalent samples (NA and HA) labeled with polyT oligo detection label (n=4)

### Assay demonstrates average accuracy of 104% recovery, average precision of 9% RSD

**Table 2** shows the results of accuracy and precision in a multi-operator study. In brief (see Methods section for details), 3 operators each analyzed 8 replicates of bivalent NA/HA in the middle of the dynamic range (24 replicates total) alongside a serial dilution of the same bivalent material used as a standard curve. Accuracy was defined as the % of the expected concentration based on the concentration back-calculated from the average standard curve, and precision shown as the % RSD of replicates. Data in **Table 2** are shown for each individual operator for the 2 capture oligos for each construct, combined over all three operators for each coding region capture oligo, and combined over all operators and all capture oligos. Single operator accuracy ranged from 97% to 110% over the 4 oligos, with an overall average of 104 (± 2) %. Single operator precision ranged from 3% to 13% RSD over the 4 oligos, with an overall average of 9 (± 2) %. No statistically significant differences were observed between operators over all capture oligos shown, as demonstrated by a two-tailed t-test (all p values were >0.14).

**Table 2.**
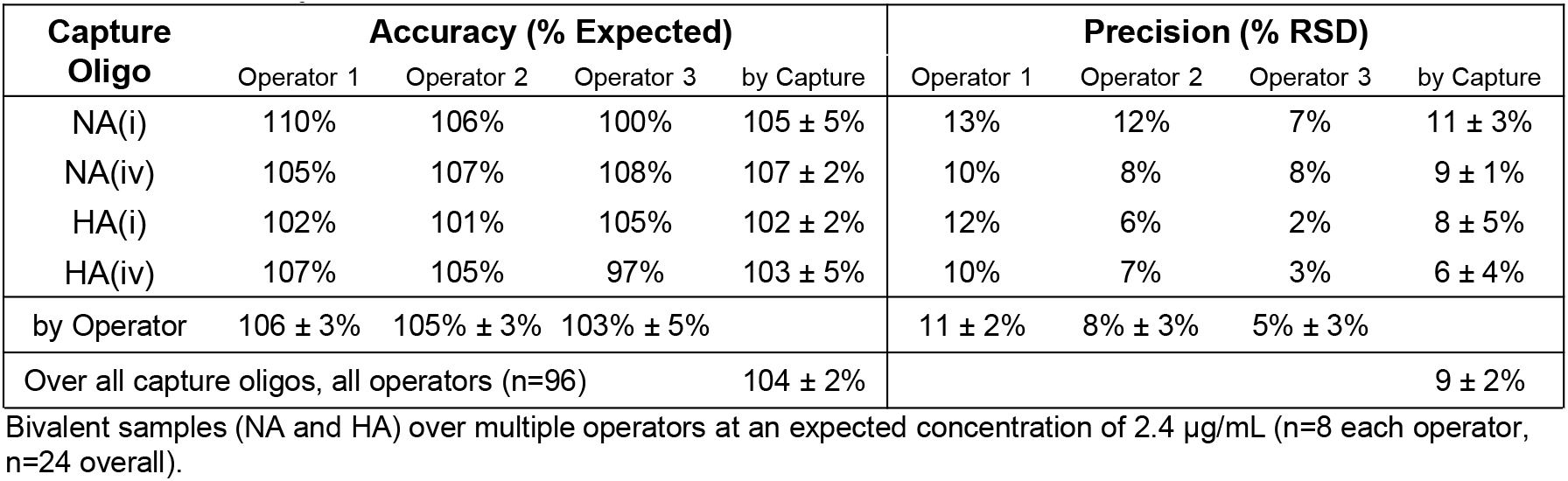
Accuracy and Precision

### Assay shows high reactivity and specificity for LNP-encapsulated mRNA

LNP encapsulation of NA and HA mRNA used the Precision NanoSystems NanoAssemblr® Spark™ microfluidic mixer following the manufacturer’s protocol, with independent quantification of encapsulated mRNA performed using the RiboGreen assay as described in the Methods section. For VaxArray analysis of LNP-encapsulated samples, samples were lysed in 1% Triton X-100 for 10 minutes after which the standard VaxArray process described was followed, with the entire assay still taking less than 2 hours. To note, no isolation (purification) of the mRNA was performed.

Blank LNPs (no mRNA encapsulated) were assessed by VaxArray post-lysis as described in the Methods section, with no non-specific signal on the microarray observed as demonstrated by S/B<1.0 (data not shown), indicating no assay cross-reactivity to the lipids themselves. **Figure 5a** compares a representative fluorescence image of the same 2 µg/mL concentration of NA mRNA in naked (unencapsulated) and LNP-encapsulated samples, both using the polyT oligo detection label and imaged at the same exposure time. These images indicate that the LNP-mRNA complex generates a similar specific response to the naked mRNA and does not produce off-target signal on the HA capture oligos. Analysis of the intensity values for each capture oligo shown in **Figure 5b** indicates good reactivity, as demonstrated by a S/B of ≥ 6.4, and specificity, as demonstrated by a S/B ≤ 1.1. Importantly, these data demonstrate good reactivity, with no requirement for upfront purification of the mRNA from LNPs, and no non-specific signal form the lipid components used as a common delivery vehicle, enabling use of the assay for formulated vaccine samples.

**Figure 5.**
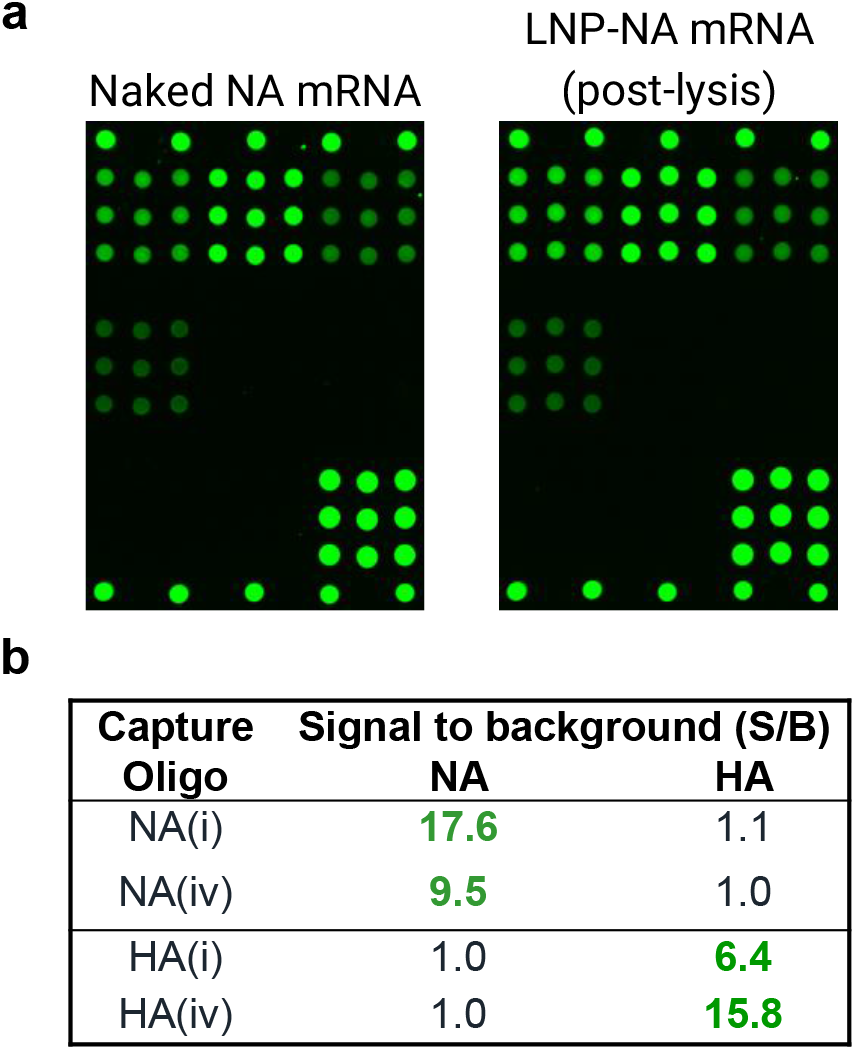
(a) Representative fluorescence images comparing naked (unencapsulated) NA mRNA and LNP-encapsulated mRNA post-lysis, both at 2 µg/mL and imaged at the same exposure time. (b) Signal/background (S/B) for monovalent LNP-encapsulated NA and HA mRNA post-lysis at 5 µg/mL. Text in bold green highlights S/B >3.

### LNP-Encapsulated mRNA shows equivalent response in mono- and bivalent samples, equivalent response to naked mRNA, and produces good accuracy and precision

Figure 6. shows a comparison of signal responses for monovalent LNP-encapsulated mRNA as well as a bivalent mixture of LNP-encapsulated NA and HA mRNA at equal concentrations in each mRNA, all labeled with the polyT oligo detection label. As shown in **Figures 6a and 6b** for the two NA-specific capture oligos, the response curves for the NA mRNA component in monovalent vs. bivalent preparations in which HA mRNA was also present are quite similar. Likewise, in **Figures 6c and 6d**, the response curves for the HA mRNA component are also similar for both the HA-specific capture oligos regardless of whether alone or in a bivalent mixture with NA mRNA. Importantly, these data show no interference from the off-target mRNA in the presence of the target mRNA and indicate independent quantification is feasible in a bivalent LNP-encapsulated mRNA mixture.

Figure 7. shows a comparison of signal responses for bivalent LNP-encapsulated mRNA as well as bivalent naked mRNA (no LNPs) using the polyT detection label, with both sets of samples processed with 1% Triton X-100 as described in the Methods section to ensure matrix matching. The response curves for both NA-specific capture oligos in **Figures 7a** and **7b** are very similar for both the naked and LNP-encapsulated samples, as are the response curves for both HA-specific capture oligos in **Figures 7c** and **7d**. Importantly, these data indicate that a naked mRNA standard should be appropriate for enabling accurate quantification in LNP-encapsulated samples over the assay linear range, provided that both are in the same matrix.

**Table 3** shows quantification accuracy and precision for LNP-encapsulated monovalent mRNA samples against a corresponding naked mRNA standard curve for the two coding region capture oligos for each mRNA as described in the Methods section. Naked mRNA standard curves along with points corresponding to the eight replicates of LNP-encapsulated mRNA analyzed are shown in **Supplementary Figure 2**. A moving 4-point linear fit was applied to each dataset as described in the Methods section, and R^2^ was >0.95 for all fits except that including the highest concentration point. Accuracy for the NA(i) and NA(iv) capture oligos were 125% and 102% of expected concentration, respectively, and HA(i) and HA(iv) produced 104% and 98% of expected, respectively. The average accuracy was 108 (± 12) %. Precision of the back-calculated concentration, expressed as the %RSD of the 8 replicates, ranged from 7% to 10% with an overall average precision of 8 (± 1) %. While NA(i) showed slightly lower accuracy than expected, the accuracy and precision data overall were generally quite similar to the data generated for the naked mRNA samples shown in **Table 2**.

**Table 3.**
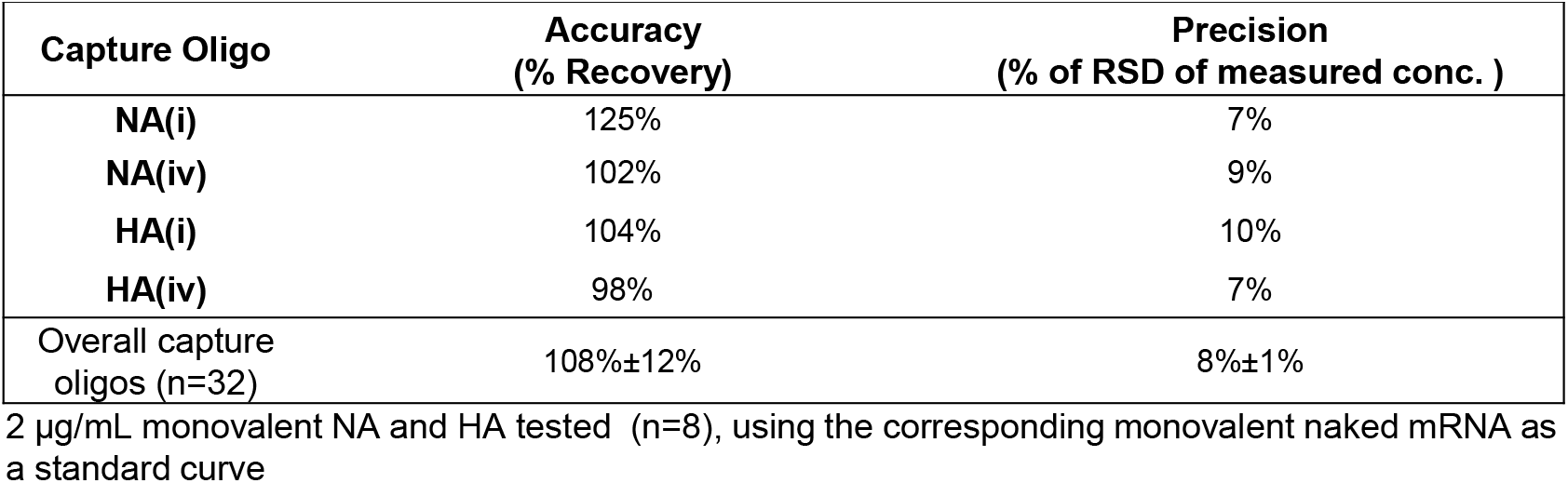
Accuracy and Precision of LNP-encapsulated mRNA

| **Capture Oligo** | **Accuracy**<br>**(% Recovery)** | **Precision**<br>**(% of RSD of measured conc. )** |
| --- | --- | --- |
| **NA(i)** | 125% | 7% |
| **NA(iv)** | 102% | 9% |
| **HA(i)** | 104% | 10% |
| **HA(iv)** | 98% | 7% |
| Overall capture<br>oligos (n=32) | 108%±12% | 8%±1% |

2 µg/mL monovalent NA and HA tested (n=8), using the corresponding monovalent naked mRNA as a standard curve

**Figure 6.**
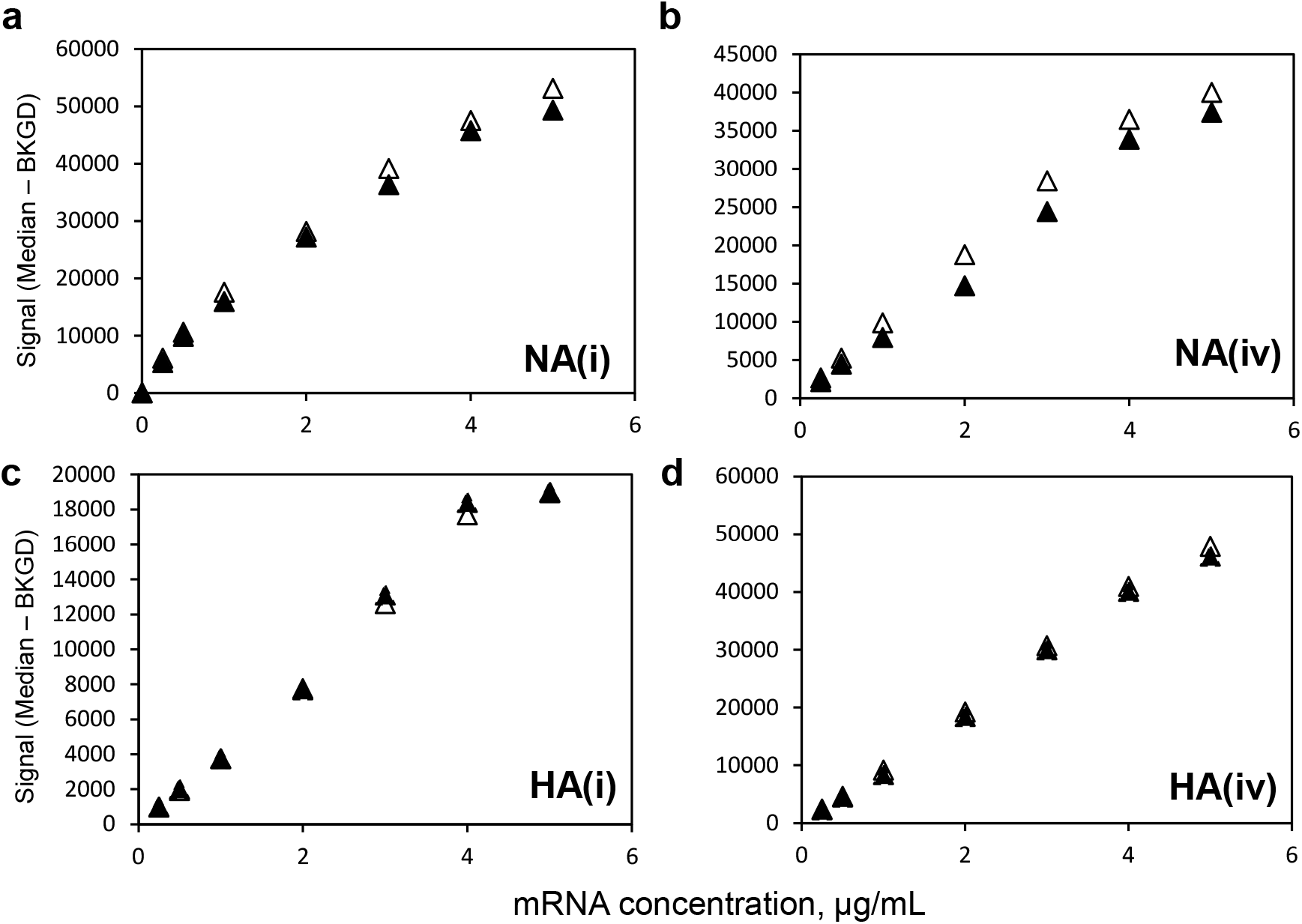
Similarity of response for monovalent (▴) and bivalent (Δ) 8-point dilution series using polyT oligo detection label for LNP-encapsulated NA mRNA using (a) NA(i) coding region capture, (b) NA(iv) coding region capture, and for LNP-encapsulated HA mRNA using (c) HA(i) coding region capture, and (d) HA(iv) coding region capture.

**Figure 7.**
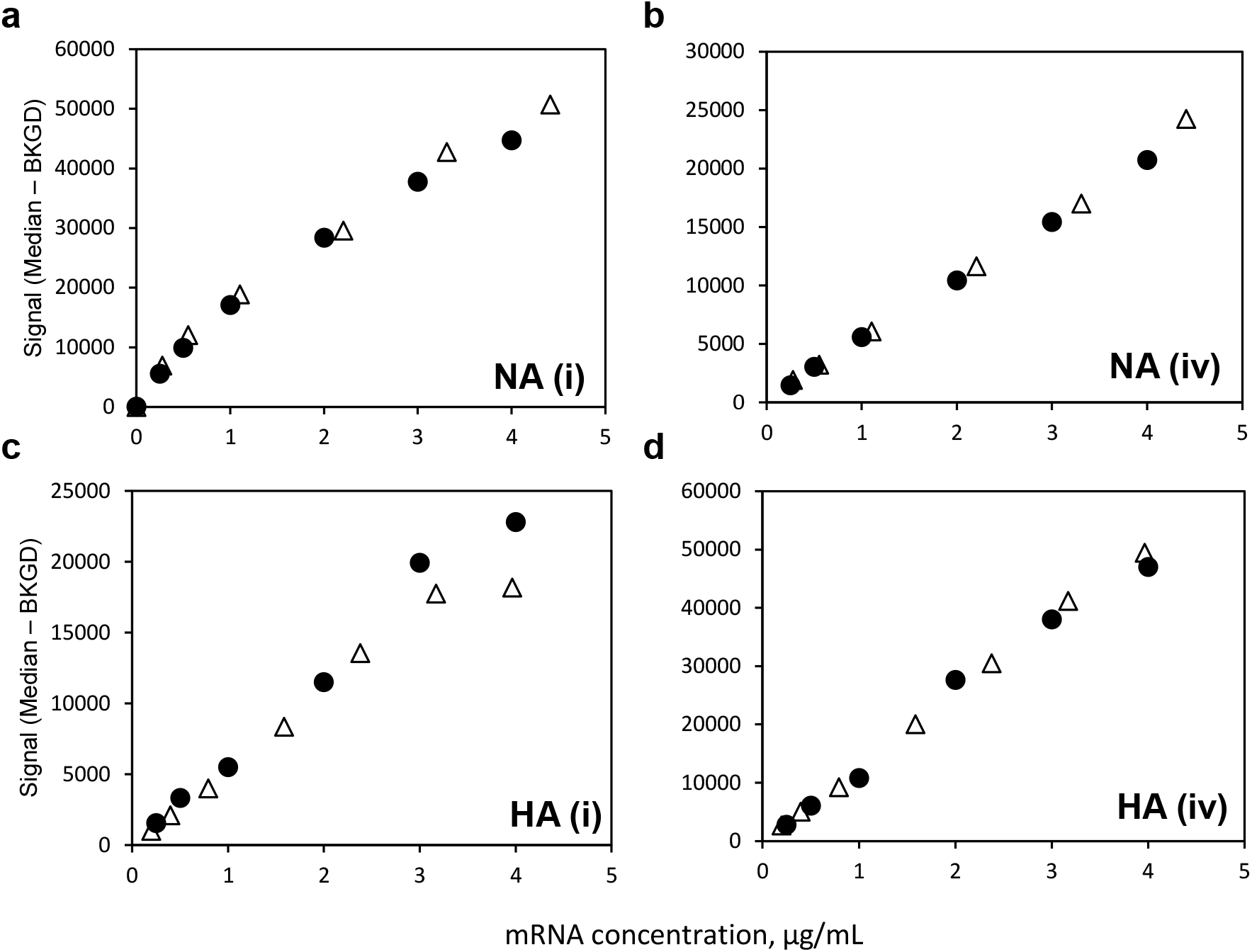
Similarity of response for bivalent LNP-encapsulated mRNA (Δ) and naked mRNA (•) 8-point dilution series using polyT oligo detection label for NA mRNA using (a) NA(i) coding region capture, (b) NA(iv) coding region capture, and for HA mRNA using (c) HA(i) coding region capture, and (d) HA(iv) coding region capture.

## DISCUSSION

With the growing field of mRNA vaccines and therapeutics and active development of numerous multivalent mRNA vaccines, it is crucial to have relevant analytical tools that can keep up with the rapid development pace and help streamline mRNA vaccine production. For use in identity and quantification applications in bioprocess development, an ideal assay should have (1) excellent analytical performance, (2) provide actionable results in a short timeframe to improve the bioprocess, (3) demonstrate good specificity to enable identity testing of multiple constructs being produced and handled at the same facility (or upon receipt after shipment between facilities), (4) have high ease of use to be run at-line and not require samples to be sent to a centralized testing laboratory to be analyzed by specially trained personnel, and (5) enable flexible throughput to enable as many or as few samples to be analyzed simultaneously as needed. In addition, for formulation development and final vaccine testing, the ideal assay should also not require additional time-consuming and complex upfront sample processing or purification steps and be free of interference from the LNP components.

The VaxArray mRNA assay described herein relies on a panel of specifically designed short capture oligos and flexible detection labeling options to enable construct-specific mRNA identity and quantification for bioprocess applications such as optimization of *in vitro* transcription and downstream purification steps. The assay provides high target specificity, critical for application to identity testing, and provides a similar level of information content as the currently recommended RT-PCR-based methods,^32^ particularly if coding region capture is paired with coding region detection labeling for added specificity. In addition, excellent linearity with dilution, dynamic range of ∼100x or more, and similar response curves in both monovalent and bivalent formulations enable quantification over relevant concentration ranges in both mono- and multivalent vaccine samples with good accuracy and precision. In addition, the assay can be uniquely used to assess quantity of full-length mRNA in monovalent formulations by capturing and labeling opposite ends of the construct. As noted, current methodologies suffer from interference in LNP-encapsulated samples, often requiring day-long upfront purification that complicates methodologies and slows down time to result.^17,32^ In LNP-encapsulated samples, the same VaxArray mRNA assay procedure used for naked mRNA samples is easily applied with only the addition of a simple 10-minute lysis step and no subsequent purification required prior to analysis. Similar analytical performance is achieved in LNP-encapsulated materials when compared to naked mRNA. The user can process and analyze a single slide (16 wells) or up to 4 slides (64 wells) simultaneously in under 2 hours, combining flexible throughput for different applications with a rapid time to result, enabling same-day actionable results.

In summary, this work demonstrates the performance of a VaxArray mRNA assay based on a nucleic acid microarray as an alternative to current methods, using a bivalent influenza mRNA system as a model for relevant multivalent vaccines currently in development. While the microarray presented here has been specifically designed to assess published influenza HA and NA constructs as a model test case, the assay can easily be optimized to target a wide variety of relevant mRNA vaccine targets in just a few months. The assay provides the benefits of a rapid time to result of under two hours, direct applicability to multivalent mRNA vaccine samples, no requirement for PCR-based amplification, and no mRNA purification needed to accurately analyze LNP-encapsulated materials relevant for formulation optimization and final mRNA vaccine formulations. These characteristics along with similar performance for both naked and LNP-encapsulated materials make this single platform attractive for identity testing and mRNA quantification even in multivalent vaccine samples throughout all stages of mRNA vaccine development, from early bioprocess development and optimization through final vaccine formulation, release, and stability.

## METHODS

### mRNA Constructs

Influenza NA and HA mRNA construct coding region sequences from Freyn et al.^35^ were synthesized commercially by Trilink Biotechnologies (San Diego, CA), including a ∼128 nt poly A tail and untranslated (UTR) regions at both the 3’ and 5’ ends. The NA construct codes for a full-length membrane-bound NA from pdm (post 2009 H1N1 A/Michigan/45/2015 (construct shown schematically in **Figure 1a**), and the shorter HA construct codes for the conserved HA stalk domain of pre-2009 H1N1 A/Brisbane/59/2007 (see **Figure 1b**). Construct sequences are listed in **Supplementary Table 1**.

### Oligonucleotide Sequence Design and Microarray Printing

Oligonucleotide capture sequences (∼20-mers) were designed to target either the NA mRNA construct (21 capture oligos) or HA mRNA construct (16 capture oligos) based on the construct sequences.^10^ Unique sequence regions for the NA and HA mRNAs anticipated to provide specificity for each were identified by first aligning the mRNA sequences in BioEdit (v7.2., (Manchester, United Kingdom). The series of 19-24-nt length sequences identified that were unique to each construct were then imported into OligoAnalyzer™ (IDT; Coralville, IA) along with the HA and NA mRNA sequences to assess sequence parameters for anticipated microarray suitability, including: self-interactions with ΔG > -7.5 kcal/mole to minimize potential for hairpin formation, absence of low complexity sequence regions (such as repeat bases of > 4nt in length), and melting temperature >52°C to enable room temperature hybridization. The ΔG was evaluated for the off-target construct to reduce potential for cross-annealing/non-specificity. Final specificity and subsequent down-selection were determined experimentally as described later. Capture sequences were ordered from IDT (Coralville, IA) at high purity. In addition, a 30-nt polyT oligonucleotide was designed to target the 3’ poly A tail of the mRNA constructs.

Oligonucleotide captures were printed at InDevR using proprietary processes onto epoxide-functionalized glass using a piezoelectric microarray printing system. An initial microarray with all 38 designed capture sequences was printed and assessed for basic reactivity and specificity for the construct of interest using the polyT oligo detection label. From this initial assessment, 4 capture oligos for each construct from a range of positions along the coding region that exhibited both high reactivity and specificity as described later were chosen for the final microarray design and printed.

### Detection Labels

Select oligonucleotides designed were synthesized by IDT with a 5’ Cy3 modification at HPLC purification grade to enable downstream fluorescence imaging using the VaxArray Imaging System. Oligonucleotides utilized as labels included the polyT oligonucleotide, NA nt1149-1168 label, and HA nt741-760 label (see sequences listed in **Supplementary Table 2)**. In addition, an anti-5’ cap antibody (MBL International Corp., Woburn, MA) was fluorescently conjugated in-house with an amine-reactive fluorescent dye antibody conjugation kit (Biotium, Fremont, CA) and used as a detection label.

### VaxArray mRNA Assay

The VaxArray mRNA nucleic acid microarray assay follows a similar overall assay protocol to previously described VaxArray assays, ^36-39^ with the slide layout, microarray layout, and a variety of possible microarray hybridization and detection schemes depicted in **Figures 1c, d**, and **e**, respectively. VaxArray slides were first equilibrated at room temperature for 30 minutes and placed inside a humidity chamber (VX-6204, InDevR Inc.). Microarray slides were pre-washed with 50 µL 1x mRNA Wash Buffer 1 (VXI-6317, InDevR Inc.) in a humidity chamber on a shaker at 80 rpm for 1 minute at 25°C. All subsequent washes and incubations were on a shaker at 80 rpm at 25°C. After the wash, samples were diluted in an optimized mRNA Oligo Binding Buffer (VXI-6316, InDevR Inc.) at a final 1x concentration and applied to designated arrays. For LNP-encapsulated mRNA samples, an additional lysis step was included prior to mixing the sample with the binding buffer in which samples were lysed in 1% Triton X-100 at 37°C for 10 minutes with all subsequent dilutions occurring in mRNA Oligo Binding Buffer supplemented with 1% Triton-X 100. Slide(s) were incubated for 1 hour and then washed with mRNA Wash Buffer 1 for 1 minute, followed by detection label incubation for 30 minutes. The slide(s) were then washed with mRNA Wash Buffer 1 once, and mRNA Wash Buffer 2 (VXI-6318, InDevR Inc.) twice prior to drying the slides by first pipetting off excess liquid from the wells followed by centrifugation for 10 s to remove any remaining liquid. Slide(s) were imaged on the VaxArray Imaging System (VX-6000, InDevR Inc.), and downstream data analysis was performed using the VaxArray Analysis Software.

### Reactivity/Specificity

Specificity of the oligos for the respective gene-specific capture oligos was verified using monovalent naked mRNA at 10 µg/mL. A label-only blank (no mRNA) was also analyzed to evaluate any direct binding of the detection labels to the capture oligos. Signal to background (S/B) ratios on all capture oligos on the microarray were calculated to assess reactivity and specificity, with a minimum reactivity threshold defined as a signal to background for the target mRNA of at least 3.0, and ideal specificity defined as a signal to background of 1.0 on off-target capture oligos.

### Analytical sensitivity and dynamic range

To estimate lower and upper limits of quantification (LLOQ and ULOQ), monovalent HA and NA mRNA constructs were combined to formulate a bivalent stock sample. From the bivalent stock, 3 replicate 16-point dilution series were prepared individually and processed as described above. The average of the 3 replicates was determined at each concentration, and a moving 4-point linear fit was applied to each dataset, calculating the slope and R^2^ for each regression.

The LLOQ was estimated by determining the concentration (within the lowest 4-point concentration range meeting minimum R^2^>0.95 requirement) at which the signal was equal to the average blank signal + 5σ (where σ = standard deviation of the background signal). To verify the LLOQ, a bivalent 8-point dilution series that spanned the approximate LLOQ for each capture oligo was created as a standard curve and analyzed alongside 8 different concentrations (n=4 at each concentration) of the bivalent sample near the approximate LLOQ. The same 4-point moving fit as described above was applied to the standard curve, and the concentrations in each replicate sample back calculated. The verified LLOQ was reported as the lowest concentration at which both the % difference from expected (accuracy) and the % relative standard deviation (RSD) (precision) of the 4 replicates were both less than 15%. ULOQ was approximated by determining the concentration at which the signal was 90% of the maximum observed signal within the previously determined concentration range. Similar to the LLOQ determination, the ULOQ was verified by evaluating 8 concentrations (n=4 each concentration) near the approximated ULOQ and determining the highest concentration at which the precision and accuracy requirements were met. Dynamic range of the assay was calculated as ULOQ/LLOQ.

### Accuracy and Precision of Naked mRNA

Three users analyzed eight (8) replicates of a contrived bivalent HA/NA mRNA sample at 2.4 µg/mL (3 users x 8 replicates = 24 replicates) using the polyT oligo detection label at 1 µM alongside an 8-point standard curve of the same contrived bivalent HA/NA sample. To investigate accuracy, the concentration in each replicate was determined by back-calculating against the fit to the average of the 3 standard curves as the expected value. Accuracy was calculated as the % of expected (measured divided by expected, expressed as a percentage), and quantified for each user individually and over the three users combined. Assay precision was measured for each user as well as over all three users combined and expressed as % RSD of replicate measurements.

### LNP Encapsulation of mRNA

The Precision Nanosystems Inc. (Vancouver, BC, Canada) NanoAssemblr® Spark™ microfluidic mixer was used to encapsulate mRNA in LNPs. Following the manufacturer’s protocol, NA and HA naked mRNA constructs were encapsulated at known quantity in the manufacturer’s provided LNP mix containing a proprietary mix of the following four lipids in ethanol: 1,2-Dioleoyl-3-trimethylammonium propane (DOTAP), distearoylphosphatidylcholine (DSPC), cholesterol, and 1,2-Dimyristoyl-*rac*-glycero-3-methoxypolyethylene glycol-2000 (PEG-DMG). Encapsulated materials were stored at +2-8°C prior to analysis.

Quantity of encapsulated mRNA and encapsulation efficiency were measured with the Quant-it™ RiboGreen RNA Assay Kit (cat# R11490; Invitrogen, Waltham, MA) following the manufacturer’s protocol. Briefly, total mRNA was measured using the RiboGreen assay with an upfront lysis step in 2% Triton X-100 (SX100-500ML; Sigma Aldrich, St. Louis, MO) to lyse the LNPs. Free mRNA was measured in the absence of the lysis step, and encapsulated mRNA was measured via subtraction of free mRNA from total mRNA, with encapsulation efficiency expressed as encapsulated mRNA divided by total mRNA, expressed as a percentage. LNP-encapsulated NA and HA mRNAs contained ∼75-85 μg/mL total mRNA, depending on the experiment, with measured encapsulation efficiencies >90%.

### Response Curves, Accuracy and Precision of Encapsulated mRNA

For assessing general response in encapsulated materials, LNP-encapsulated monovalent NA and HA mRNA and a bivalent mixture of NA and HA mRNA were analyzed to assess similarity of responses. LNP-encapsulated bivalent NA and HA mRNA and a bivalent mixture of naked mRNA were compared to assess similarity of responses. For accuracy and precision measurements, monovalent NA and HA mRNA were separately encapsulated in LNPs and analyzed post-lysis in 8 replicates alongside a standard curve of naked mRNA also pre-treated in 1% Triton X-100 to match the sample matrix. Replicates were prepared at 2 µg/mL total mRNA. Signals from the lysed replicates were back-calculated using the standard curve to determine the measured concentration and associated accuracy and precision. All samples were run in 1x mRNA oligo binding buffer with 1% Triton X-100 to ensure matrix matching.

## DATA AVAILABILITY STATEMENT

All relevant data from this study are available from the authors.

## ACKNOWLEDGEMENTS

We sincerely thank the laboratory of Jay Hesselberth, School of Medicine, University of Colorado Anschutz Medical Campus, for access to the NanoAssemblr® Spark™ instrument.

## AUTHOR CONTRIBUTIONS

R.G. supervision, methodology, investigation, validation, formal analysis, visualization, writing-original draft, writing-review and editing; C.R. methodology, investigation, validation, formal analysis, visualization, writing-original draft, writing-review and editing; E.T. oligo capture design, visualization, writing-review and editing; R.B. visualization, writing-review and editing; M.G. methodology, investigation, validation, writing-one section methods, writing-review and editing; C.M. methodology, investigation, formal analysis, writing-review and editing; A.T. project administration, resources, visualization, writing-review and editing; T.H. resources, visualization, writing-review and editing; K.R. conceptualization, project administration, resources, writing-review and editing; E.D. project administration, supervision, methodology, resources, formal analysis, writing-original draft, writing-review and editing, visualization.

## SUPPLEMENTARY FIGURE LEGENDS

**Supplementary Figure 1.** Linearity of response for a variety of capture and detection schemes in monovalent mRNA samples, with NA in left column and HA in right column. a) Coding region capture with polyT label as shown in Figure 1e(i), (b) Coding region capture and labeling with 5’ cap antibody label as shown in Figure 1e(ii). Data point are the average of three replicates and error bars indicate ± 1 standard deviation (n=3). R^2^ in upper left of each plot is based on a single linear regression.

**Supplementary Figure 2.** Signal Response curves for monovalent LNP-mRNA samples, labeled with polyT label. 8-pt response curves generated comparing a monovalent standard curve (•) to eight replicates of lysed LNP-mRNA (Δ) using (a) NA(i) coding region capture oligo, (b) NA(iv) coding region capture oligo, and for HA mRNA using (c) HA(i) coding region capture oligo, and (d) HA(iv) coding region capture oligo.

