## Supplemental Figures and Tables for "Rapid Identity and Quantity CQA Test for Multivalent mRNA Drug Product Formulations"

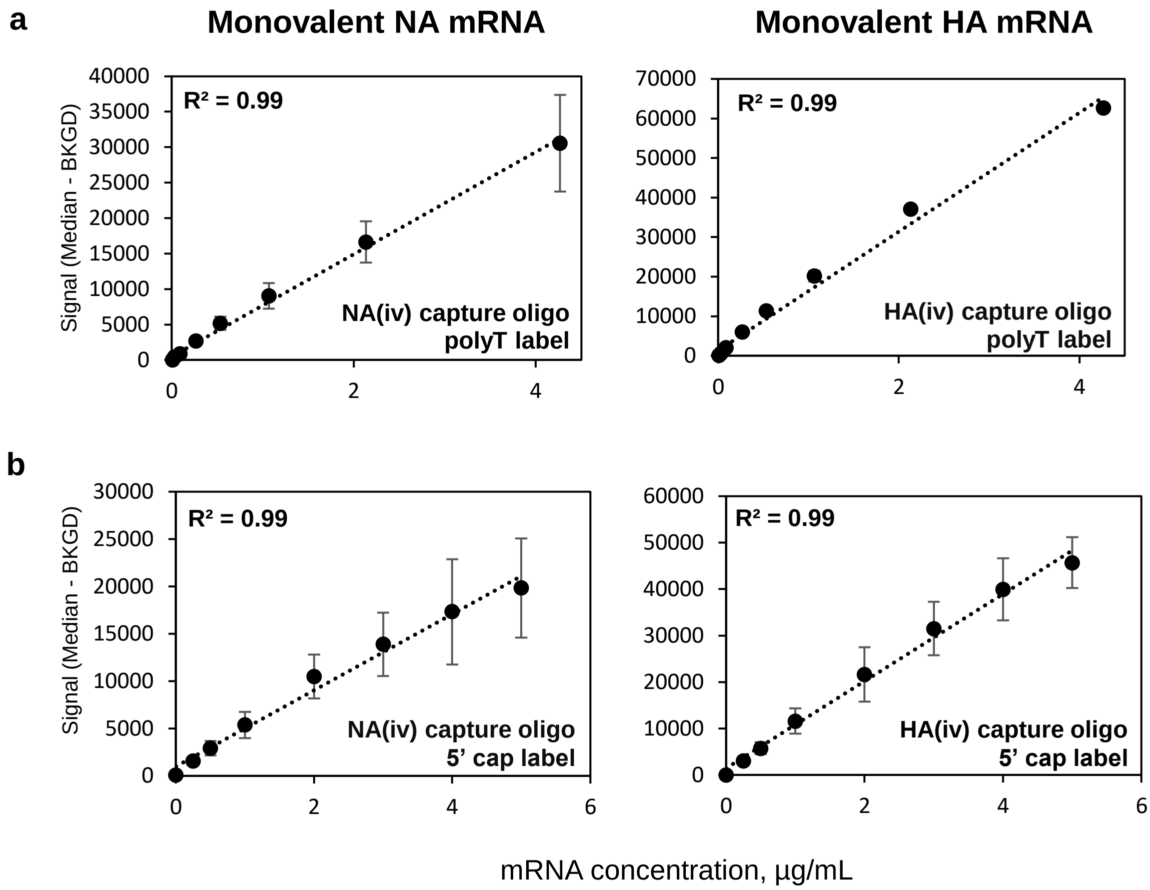


**Supplementary Figure 1.** Linearity of response for a variety of capture and detection schemes in monovalent mRNA samples, with NA in left column and HA in right column. a) Coding region capture with polyT label as shown in Figure 1e(i), (b) Coding region capture and labeling with 5' cap antibody label as shown in Figure 1e(ii). Data point are the average of three replicates and error bars indicate ± 1 standard deviation (n=3). $R^2$ in upper left of each plot is based on a single linear regression.

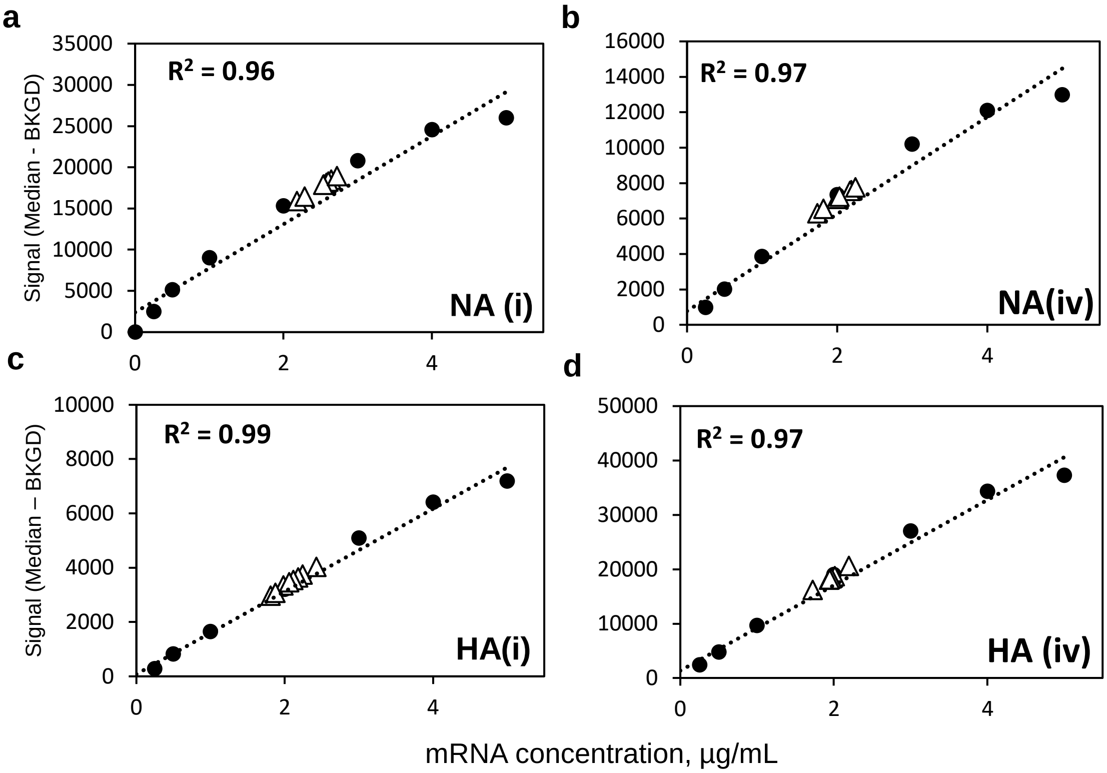


**Supplementary Figure 2.** Signal Response curves for monovalent LNP-mRNA samples, labeled with polyT label. 8-pt response curves generated comparing a monovalent standard curve (●) to eight replicates of lysed LNP-mRNA (Δ) using (a) NA(i) coding region capture oligo, (b) NA(iv) coding region capture oligo, and for HA mRNA using (c) HA(i) coding region capture oligo, and (d) HA(iv) coding region capture oligo.

**Supplementary Table 1**. NA and HA construct sequences

| mRNA | Sequence |
|---|---|
| **NA** | ATGAACCCCAACCAGAAGATCATCACCATCGGCTCCATCTGCATGACCATCGGCATGGCCAACCTGATC<br>CTGCAGATCGGCAACATCATCTCCATCTGGGTGTCCCACTCCATCCAGATCGGCAACCAGTCCCAGATC<br>GAGACCTGCAACCAGTCCGTGATCACCTACGAGAACAACACCTGGGTGAACCAGACCTACGTGAACAT<br>CTCCAACACCAACTTCGCCGCCGGCCAGTCCGTGGTGTCCGTGAAGCTGGCCGGCAACTCCTCCCTGT<br>GCCCCGTGTCCGGCTGGGCCATCTACTCCAAGGACAACTCCGTGCGCATCGGCTCCAAGGGCGACGT<br>GTTCGTGATCCGCGAGCCCTTCATCTCCTGCTCCCCCCTGGAGTGCCGCACCTTCTTCCTGACCCAGG<br>GCGCCCTGCTGAACGACAAGCACTCCAACGGCACCATCAAGGACCGCTCCCCCTACCGCACCCTGATG<br>TCCTGCCCCATCGGCGAGGTGCCCTCCCCCTACAACTCCCGCTTCGAGTCCGTGGCCTGGTCCGCCTC<br>CGCCTGCCACGACGGCATCAACTGGCTGACCATCGGCATCTCCGGCCCCGACTCCGGCGCCGTGGCC<br>GTGCTGAAGTACAACGGCATCATCACCGACACCATCAAGTCCTGGCGCAACAACATCCTGCGCACCCA<br>GGAGTCCGAGTGCGCCTGCGTGAACGGCTCCTGCTTCACCATCATGACCGACGGCCCCTCCGACGGC<br>CAGGCCTCCTACAAGATCTTCCGCATCGAGAAGGGCAAGATCATCAAGTCCGTGGAGATGAAGGCCCC<br>CAACTACCACTACGAGGAGTGCTCCTGCTACCCCGACTCCTCCGAGATCACCTGCGTGTGCCGCGACA<br>ACTGGCACGGCTCCAACCGCCCCTGGGTGTCCTTCAACCAGAACCTGGAGTACCAGATGGGCTACATC<br>TGCTCCGGCGTGTTCGGCGACAACCCCCGCCCCAACGACAAGACCGGCTCCTGCGGCCCCGTGTCCT<br>CCAACGGCGCCAACGGCGTGAAGGGCTTCTCCTTCAAGTACGGCAACGGCGTGTGGATCGGCCGCAC<br>CAAGTCCATCTCCTCCCGCAAGGGCTTCGAGATGATCTGGGACCCCAACGGCTGGACCGGCACCGACA<br>ACAAGTTCTCCATCAAGCAGGACATCGTGGGCATCAACGAGTGGTCCGGCTACTCCGGCTCCTTCGTG<br>CAGCACCCCGAGCTGACCGGCCTGGACTGCATCCGCCCCTGCTTCTGGGTGGAGCTGATCCGCGGCC<br>GCCCCGAGGAGAACACCATCTGGACCTCCGGCTCCTCCATCTCCTTCTGCGGCGTGAACTCCGACACC<br>GTGGGCTGGTCCTGGCCCGACGGCGCCGAGCTGCCCTTCACCATCGACAAG |
| **HA** | ATGAAGGTGAAGCTGCTGGTGCTGCTGTGCACCTTCACCGCCACCTACGCCGACACCATCTGCATCGG<br>CTACCACGCCAACAACTCCACCGACACCGTGGACACCGTGCTGGAGAAGAACGTGACCGTGACCCACT<br>CCGTGAACCTGCTGGAGAACGGCGGCGGCGGCAAGTACGTGTGCTCCGCCAAGCTGCGCATGGTGAC<br>CGGCCTGCGCAACAAGCCCTCCAAGCAGTCCCAGGGCCTGTTCGGCGCCATCGCCGGCTTCACCGAG<br>GGCGGCTGGACCGGCATGGTGGACGGCTGGTACGGCTACCACCACCAGAACGAGCAGGGCTCCGGC<br>TACGCCGCCGACCAGAAGTCCACCCAGAACGCCATCAACGGCATCACCAACAAGGTGAACTCCGTGAT<br>CGAGAAGATGAACACCCAGTACACCGCCATCGGCTGCGAGTACAACAAGTCCGAGCGgTGCATGAAGC<br>AGATCGAGGACAAGATCGAGGAGATCGAGTCCAAGATCTGGTGCTACAACGCCGAGCTGCTGGTGCTG<br>CTGGAGAACGAGCGCACCCTGGACTTCCACGACTCCAACGTGAAGAACCTGTACGAGAAGGTGAAGTC<br>CCAGCTGAAGAACAACGCCAAGGAGATCGGCAACGGCTGCTTCGAGTTCTACCACAAGTGCAACGACG<br>AGTGCATGGAGTCCGTGAAGAACGGCACCTACGACTACCCCAAGTACTCCGAGGAGTCCAAGCTGAAC<br>CGCGAGAAGATCGACGGCGTGAAGCTGGAGTCCATGGGCGTGTACCAGATCGAGGGCCGC |

**Supplementary Table 2**. Oligonucleotide sequences designed

| Type | Name | Sequence (5' → 3') |
|---|---|---|
| Capture oligo | NA(i) | GATGTTGCCGATCTGCAGGATC |
| | NA(ii) | GTTGTTGCGCCAGGACTTGAT |
| | NA(iii) | GTTCTGGTTGAAGGACACCCAGG |
| | NA(iv) | TGATGGAGAACTTGTTGTCG |
| | HA(i) | TGTTGGCGTGGTAGCCGATGC |
| | HA(ii) | CCGCTCGGACTTGTTGTACTC |
| | HA(iii) | TGCACTCGTCGTTGCACTTGTG |
| | HA(iv) | CGTCGATCTTCTCGCGGTTC |
| | polyT | TTTTTTTTTTTTTTTTTTTTTTTTTTTTTT |
| Label | Poly T | TTTTTTTTTTTTTTTTTTTTTTTTTTTTTT |
| | NA nt1149-1168 | TGATGGAGAACTTGTTGTCG |
| | HA nt741-760 | CGTCGATCTTCTCGCGGTTC |
